# A theoretical framework for the site-specific and frequency-dependent neuronal effects of deep brain stimulation

**DOI:** 10.1101/2020.11.30.404269

**Authors:** Luka Milosevic, Suneil K Kalia, Mojgan Hodaie, Andres M Lozano, Milos R Popovic, William D Hutchison, Milad Lankarany

## Abstract

With the growing interest in the expansion of deep brain stimulation indications, we aimed to provide experimental and computational insights into the brain-region-specific and frequency-dependent effects of extracellular stimulation on human neuronal activity. Experimentally, we demonstrated microstimulation-evoked excitatory neuronal responses in the ventral intermediate nucleus and reticular thalamus, and inhibitory responses in the subthalamic nucleus and substantia nigra pars reticulata; hypothesized to be the result of simultaneous activations of convergent afferent inputs. Higher stimulation frequencies led to a loss of site-specificity and convergence towards neuronal suppression; hypothesized to be mediated by synaptic depression. These experimental findings were reproduced by a computational framework in which relative distributions of convergent excitatory/inhibitory afferents were embedded within a model of short-term synaptic plasticity for the prediction of site-specific and frequency-dependent responses to extracellular stimulation. This theoretical framework may aid in the design of physiologically-informed stimulation paradigms in existing or prospective deep brain stimulation indications.

## Introduction

Deep brain stimulation (DBS) is an established neuromodulatory therapy for several movement disorders including Parkinson’s disease (Limousin et al., 1998), essential tremor (Dallapiazza et al., 2019), and dystonia (Hung et al., 2007). DBS has also recently received approvals for the treatment of obsessive-compulsive disorder (Menchón et al., 2019) and epilepsy (Fisher et al., 2010). Despite a rapidly growing interest in the development of new DBS indications (Youngerman et al., 2016), the various ways in which DBS may influence neuronal activity are not fully understood. In addition to appropriate patient selection, physiologically-informed target selection and subsequent stimulation programming play important roles in the development of novel indications (Brocker et al., 2017). As such, it must be considered that the effects of electrical stimulation in one area of the brain may differ from the effects of stimulation delivery in another (Basu et al., 2019). The objective of this study was to demonstrate how the neuronal effects of DBS vary depending on the stimulation target region and the frequency of stimulation pulses being delivered. Knowledge of the site-specific abilities to selectively upregulate or downregulate neuronal output is of importance for the development of physiologically-informed stimulation paradigms in existing or prospective DBS indications and could also aid in otherwise empirically-performed stimulation programming procedures.

It was previously demonstrated that single pulses of electrical stimulation delivered to the substantia nigra pars reticulata (SNr) or globus pallidus internus (GPi) were associated with stimulation-evoked inhibitory responses, likely mediated by local GABA release (Dostrovsky et al., 2000; Liu et al., 2012; Milosevic et al., 2018a); whereas high-frequency stimulation (HFS) of the thalamic ventral intermediate nucleus (Vim) elicited brief short-latency excitatory responses, likely the result of unsustained local glutamate release (Milosevic et al., 2018b). In the current study, microelectrode recordings of single-neuron activity across four brain regions (Vim, thalamic reticular nucleus (Rt), subthalamic nucleus (STN), and SNr) were assessed during microstimulation trains across a range of frequencies (1-100Hz). We hypothesized that (i) the effects of individual electrical stimulation pulses would vary with respect to the distribution of afferent inputs converging on target neurons (whether predominantly inhibitory or excitatory), and that based on these local neuroanatomical properties, stimulation pulses would elicit either net inhibitory or excitatory responses. Moreover, based on previous findings of HFS-induced depression of stimulus-evoked field potentials (Liu et al., 2012; Milosevic et al., 2018a; Steiner et al., 2019), we hypothesized that (ii) suppression of neuronal activity during HFS is mediated by changes to short-term synaptic dynamics; namely, non-selective synaptic depression (i.e. depression of both inhibitory and excitatory synaptic transmission). Indeed, experimental work in rodent STN slices has demonstrated that pharmacologically-isolated excitatory and inhibitory postsynaptic potentials were both depressed during HFS (Steiner et al., 2019).

In addition to experimental data collection, we developed a computational framework for the prediction of site-specific and frequency-dependent neuronal responses to electrical stimulation based on the above hypotheses. Previous theoretical works suggest that individual pulses of extracellular stimulation (i.e. DBS) initiate action potentials which are propagated along the axons of stimulated neurons (Anderson et al., 2018); this includes both efferent axons and afferent axons and/or their terminals (Jakobs et al., 2019; McIntyre et al., 2004). These axonal activations can in turn mediate synaptic transmission. To this end, it was shown that HFS may reduce synaptic transmission fidelity by way of synaptic depression (Farokhniaee and McIntyre, 2019) or axonal failure (Rosenbaum et al., 2014). What has not typically been considered in previous computational work is that anatomical differences across stimulation sites may incur dissimilar postsynaptic responses (i.e. neuronal excitation vs. inhibition). Based on our first hypothesis, our computational model considers that the postsynaptic neuronal responses to individual DBS pulses are the result of a simultaneous activation of presynaptic inputs and takes into consideration the site-specific proportions of inhibitory and excitatory inputs converging on target neurons (derived from anatomical/ morphological literature). In accordance with our second hypothesis, the Tsodyks-Markram (TM) model (Tsodyks and Markram, 1997) of short-term synaptic plasticity was embedded within our computational framework in order to account for changes to synaptic transmission fidelity based on the frequency of successive stimuli. Despite the relative success of previous models in explaining the mechanisms of high-frequency DBS amongst homogenous cell-types, to the best of our knowledge, no theory has addressed the biophysical mechanisms underlying DBS at different frequencies across various brain regions within a single computational framework; especially within the context of the *in vivo* human brain.

## Results

### Experimental: Responses to single stimulation pulses

The responses to single stimulation pulses (Fig. 1A) revealed stimulus-evoked excitatory responses for Vim and Rt, and inhibitory responses for STN and SNr. For Vim, the average firing rates of the immediate 20ms (p=.002) and 40ms (p=.003) periods following stimulation pulses were significantly greater than the 20ms pre-stimulus period. This was also the case for the 20ms (p<.001) and 40ms (p<.001) post-stimulus periods for Rt. For STN, the average firing rates of the 20ms (p<.0001) and 40ms (p=.003) post-stimulus periods were significantly less than the 20ms pre-stimulus period. This was also the case for the 20ms (p<.0001) and 40ms (p<.001) post-stimulus periods for SNr. All statistics have been corrected for multiple comparisons. Cohen’s d_z_ effect sizes are depicted in Fig. 1A.

**Fig. 1.**
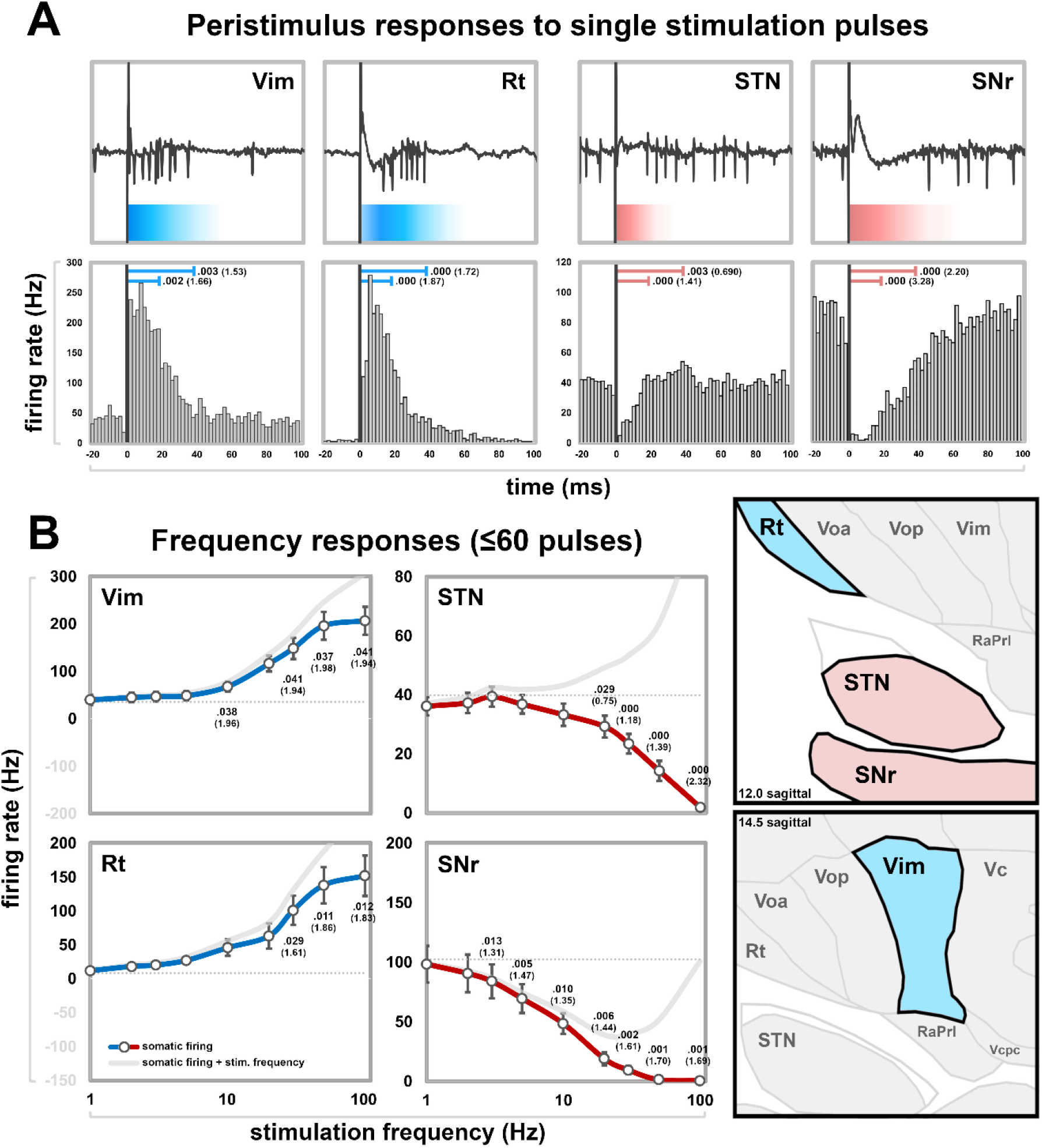
Experimental (A) peristimulus responses and (B) frequency response functions. (A) Top panels show exemplary responses to a single stimulation pulse in each structure, whereas bottom panels show groupwise peristimulus time histograms of stimulus-evoked excitatory responses for Vim and Rt and stimulus-evoked inhibitory responses for STN and SNr. The average firing rates of the immediate 20ms and 40ms periods following stimulations pulses were significantly greater than the 20ms pre-stimulus periods for Vim and Rt, and significantly lesser for STN and SNr (Bonferonni-corrected 2-tailed paired t-tests, Cohen’s d_z_ effect sizes in parentheses). (B) Stimulation (≤60 pulses) frequency response functions show that average firing rates progressively increased in Vim and Rt as the stimulation frequency became greater, while they progressively decreased in STN and SNr. The average ± standard error baseline firing rates for Vim, Rt, STN, and SNr neurons were 32.0±11Hz, 8.2±1Hz, 39.9±3Hz, and 102.3±16Hz, respectively (dashed gray lines). Firing rates during the various stimulation trains were compared to the baseline firing rates and the results of Bonferroni-corrected post hoc t-tests (2-tailed, paired) are displayed with Cohen’s d_z_ effect sizes in parentheses. In the case that each DBS pulse generates an action potential on the efferent axon, then the overall neuronal output would be the summation of the somatic firing rate and stimulation frequency; this is represented by the solid gray lines in each plot. The right anatomical panels are 12.0mm and 14.5mm sagittal sections.

### Experimental: Stimulation frequency response functions

The average ± standard error baseline firing rates for Vim, Rt, STN, and SNr neurons were 32.0±11Hz, 8.2±1Hz, 39.9±3Hz, and 102.3±16Hz, respectively. The stimulation frequency response functions (Fig. 1B) show excitatory responses for Vim and Rt, and inhibitory responses for STN and SNr. For Vim, neuronal firing rates progressively increased as the stimulation frequency became greater and a significant main effect of stimulation was found [F=43.074 (9,234), p<.001, η^2^=0.624]. Bonferroni-corrected t-tests revealed differences in neuronal firing compared to baseline at stimulation frequencies of 10Hz (p=.038), 30Hz (p=.041), and greater (p<0.05). For Rt, neuronal firing rates also progressively increased as the stimulation frequency became greater and a significant main effect of stimulation was found [F=31.170 (9,117), p<.001, η^2^=0.706]. Statistically significant differences in neuronal firing compared to baseline were found at stimulation frequencies of 30Hz (p=.029) and greater (p<.05). For STN, neuronal firing rates were progressively attenuated as the stimulation frequency became greater and a significant main effect of stimulation was found [F=26.420 (9,91), p<.001, η^2^=0.746]. Statistically significant differences in neuronal firing compared to baseline were found at stimulation frequencies of 20Hz (p=.029) and greater (p<.001). For SNr, neuronal firing rates also progressively attenuated as the stimulation frequency became greater and a significant main effect of stimulation was found [F=25.890 (9,63), p<.001, η^2^=0.787]. Statistically significant differences in neuronal firing compared to baseline were found at stimulation frequencies of 3Hz (p<.05) and greater (p≤.01). Detailed post hoc t-test statistics (all corrected for multiple comparisons within the text and figures) and Cohen’s d_z_ effect sizes are depicted in Fig. 1B.

### Experimental: Time-domain responses to stimulation

In Vim and Rt, periodic excitatory responses were evident at 5Hz and 10Hz (Fig. 2). The strength of the excitatory responses attenuated over time during stimulation trains of ≥20Hz and were modelled by double exponential decay functions (R^2^ values within Fig. 2). The time-series histograms for long train ≥100Hz data (≥2s) in Vim and Rt show particularly prominent time-varying responses. While these stimulations did elicit excitatory responses, they were transient in nature and limited to start of stimulation. In Vim, the initial excitatory response at 200Hz was of shorter duration than at 100Hz, and the subsequent neuronal suppressive response was stronger at 200Hz than at 100Hz, with firing rates dropping below baseline. In SNr, periodic inhibitory responses were evident at 5Hz and 10Hz. In STN and SNr, there was an overall stationary neuronal suppressive effect with increasing frequency (rather than an effect which changed dynamically over time as was the case in Vim and Rt).

**Fig. 2.**
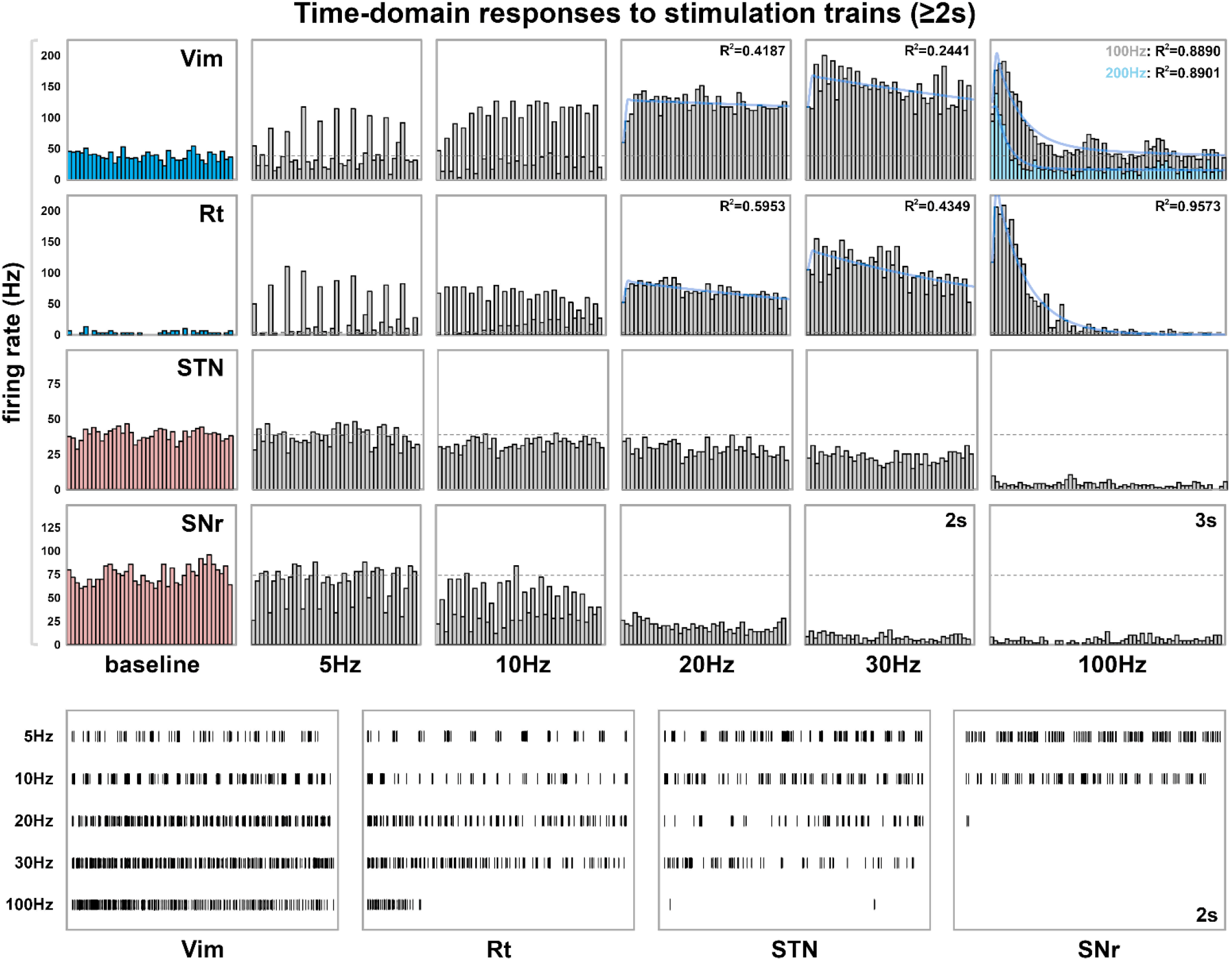
Experimental time-domain responses to stimulation trains. For Vim and Rt, firing rates progressively increased above baseline (gray dashed lines) with increasing stimulation frequencies. Periodic excitatory responses are shown at 5Hz and 10Hz, however neuronal excitation declined over time with ≥20Hz. Excitatory responses with 100Hz long trains (≥2s) were transient, and a subsequent reduction of neuronal firing is evident after the initial excitation. In Vim, the initial excitatory response at 200Hz is of shorter duration than at 100Hz, and the subsequent neuronal suppressive response is stronger at 200Hz compared to 100Hz, with firing rates dropping below baseline. In STN and SNr, firing rates progressively decreased with increasing stimulation frequencies. In SNr, periodic inhibitory responses are visible at 5 and 10Hz. Exemplary firing rate raster data from each structure during various stimulation trains are displayed in the four bottom-most panels.

### Computational: Responses to single stimulation pulses

The net changes to postsynaptic currents in response to single pulses of stimulation were modelled by simultaneous activations of all presynaptic inputs (Fig. 3Bi). These responses differed across brain regions due to differences in the proportions of excitatory and inhibitory inputs (summarized in the Methods subsection “Computational: Presynaptic inputs” and Supplementary Table 1). The simulated peristimulus firing rate histograms (i.e. the neuronal responses to the aforementioned changes to presynaptic currents) revealed stimulus-evoked excitatory responses for Vim (peak firing rate of 405.9Hz vs. 245.1Hz in the experimental data), inhibitory responses for SNr (minimum firing rate of 0Hz vs. 0.7Hz in the experimental data), and a short-latency excitatory responses (78.8Hz peak vs. no peak in the experimental data) followed by a longer latency inhibitory response (8.4Hz trough vs. 4.8Hz in the experimental data) for STN (Fig. 3Bii). The lack of short-latency excitation in the experimental data for STN might be explained by discrepancies in the temporal dynamics of excitatory transmission and/or occlusion of the excitatory response by the stimulus artifact. Figure generation for model Rt neurons was omitted due to redundancy as the model parameters are identical to Vim except for the parameter which underlies the baseline firing rate (elaborated upon in the Methods secion; model parameters for Rt are nevertheless provided in the Supplementary Material).

**Fig. 3.**
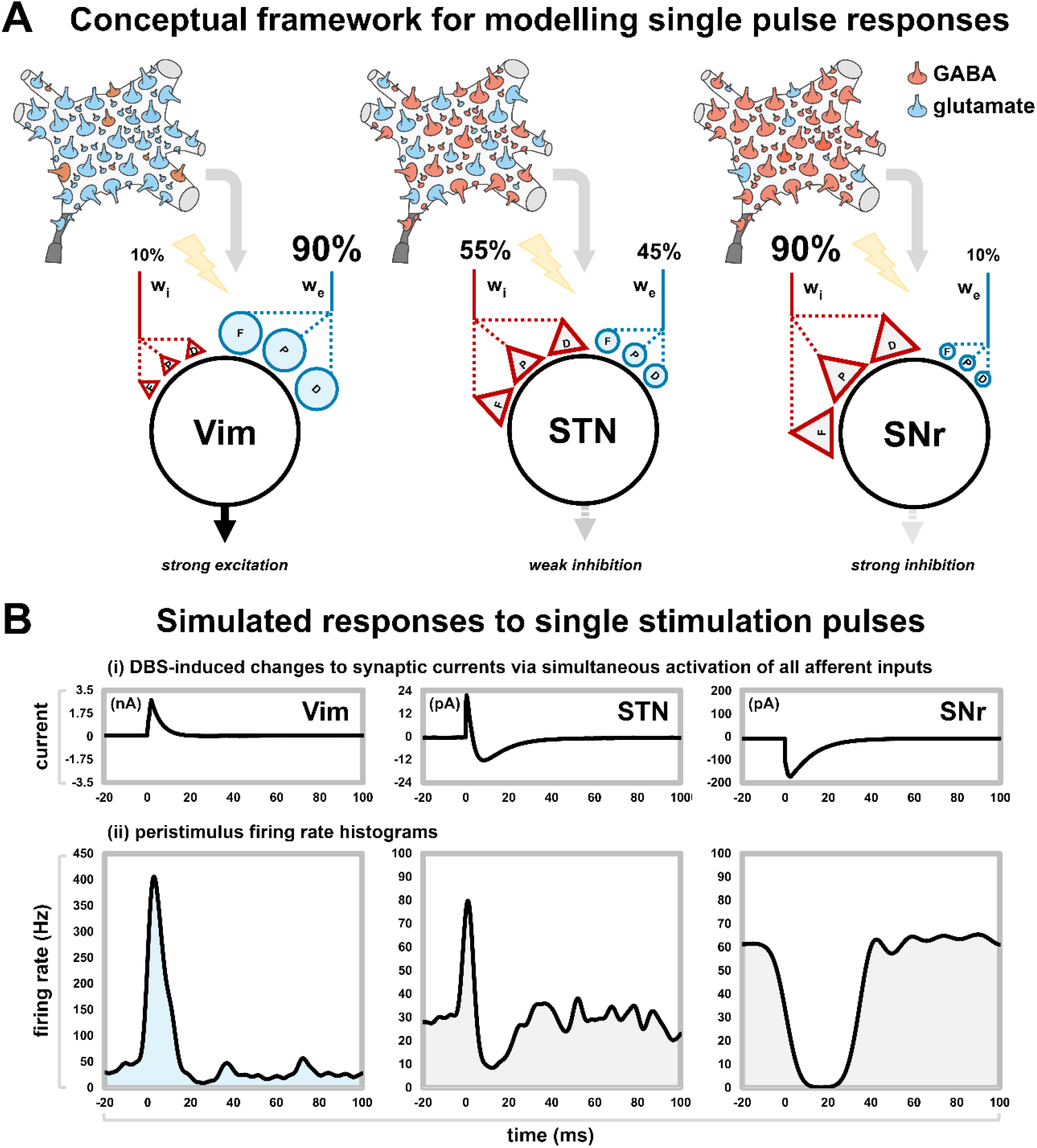
Computational (A) model framework and (B) simulated peristimulus responses. (A) To model the response to single pulses of electrical stimulation, each model neuron was assigned a certain proportion of excitatory and inhibitory presynaptic inputs/weights with proportions derived from anatomical literature. The effect of each DBS single pulse was modelled by simultaneously activating all presynaptic inputs. (B) The corresponding changes to (i) synaptic currents and (ii) somatic firing induced by the simultaneous activations are displayed (i.e. the single-pulse responses). This framework closely replicated the robust stimulus-evoked neuronal excitation in Vim and neuronal inhibition in SNr. In STN, there was a short-latency neuronal excitation which was not observed in the experimental data (though may have been occluded by the stimulation artifact) due to the high speed of excitatory synaptic transmission, followed by an inhibitory period congruent with the experimental data.

### Computational: Time-domain synaptic currents

Excitatory and inhibitory synaptic currents were generated separately, along with the total (i.e. sum of excitatory and inhibitory) synaptic currents in responses to DBS pulses across a range of frequencies for each of Vim, STN, and SNr (Fig. 4). The TM model accounts for frequency-dependent changes to short-term synaptic dynamics. In all structures, the model suggests non-selective frequency-dependent depression of both excitatory and inhibitory synaptic currents. For Vim, sustained periodic excitations are seen with 5Hz and 10Hz, while frequency-dependent weakening of the excitatory responses with successive stimuli are observed with frequencies ≥20Hz. Predominant inhibitory synaptic currents corroborate the strong inhibitions of somatic firing in SNr with low stimulation frequencies; whereas neuronal suppression with higher frequencies is likely the result of non-selective frequency-dependent synaptic depression. For STN, the mixed excitatory-inhibitory stimulus-evoked responses likely explain the more net-neutral somatic firing responses in experimental data with lower stimulation frequencies; whereas non-selective synaptic depression can explain the frequency-dependent suppression of somatic firing with higher stimulation frequencies.

**Fig. 4.**
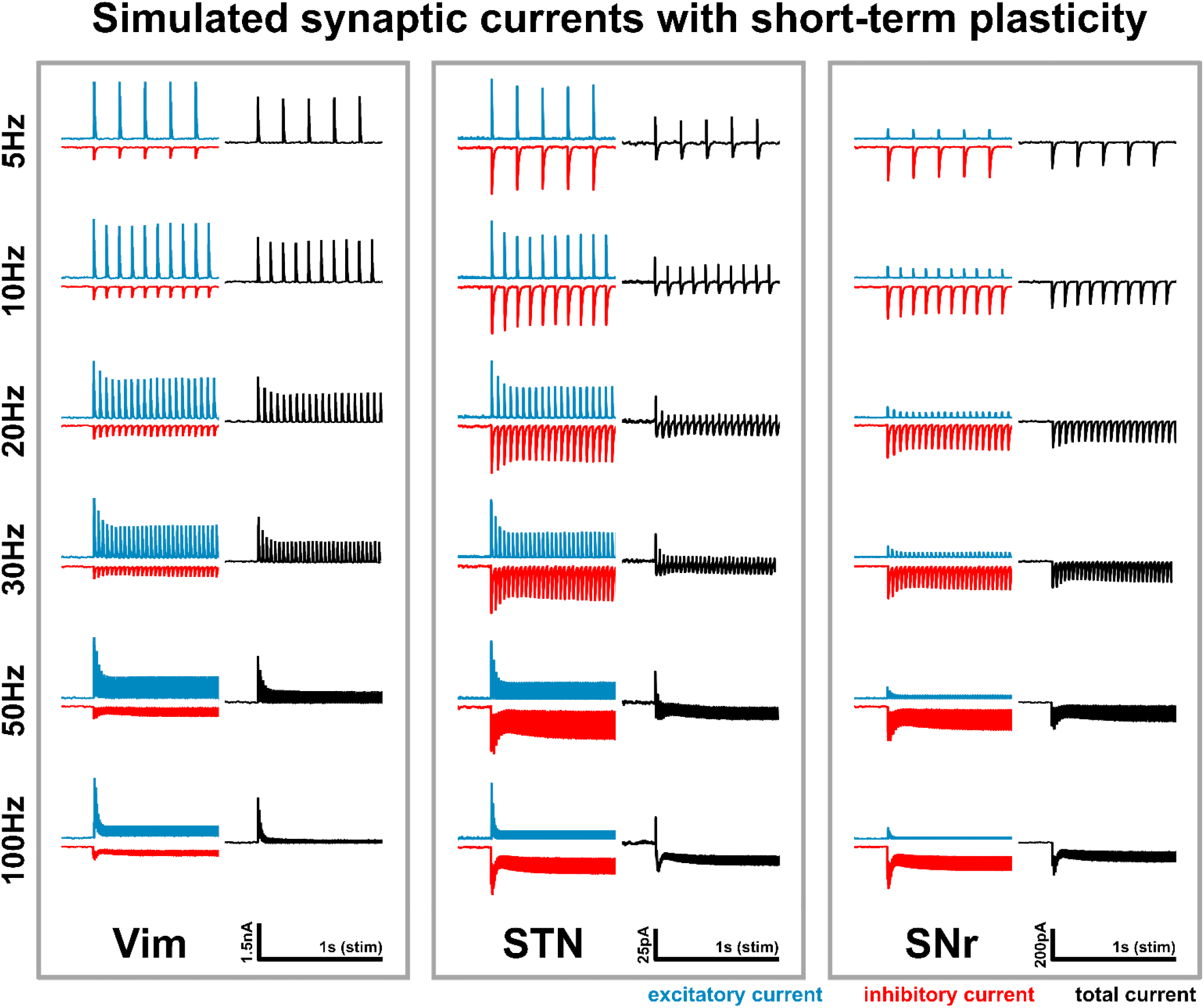
Computational time-domain synaptic currents. The three figures show excitatory, inhibitory, and total (i.e. sum of excitatory and inhibitory) synaptic currents in responses to DBS pulses across a range of frequencies with an embedded TM model to account for short-term synaptic dynamics. In all cases, the model suggests non-selective frequency-dependent depression of both excitatory and inhibitory synaptic currents. For Vim, rather stable periodic excitations are seen with 5Hz and 10Hz. Also corroborating experimental data, frequency-dependent weakening of the excitatory responses is observed with frequencies ≥20Hz. Predominant inhibitory synaptic currents corroborate the strong inhibitions of somatic firing in SNr, together with non-selective frequency-dependent synaptic depression. For STN, the mixed excitatory-inhibitory stimulus-evoked responses likely explain the more net-neutral somatic firing responses in experimental data with lower stimulation frequencies, while non-selective synaptic depression can explain frequency-dependent suppression of somatic firing.

### Computational: Time-domain membrane potentials

The membrane potentials of modelled neurons in response to DBS across a range of frequencies were generated for each of Vim (Fig. 5), STN (Fig. 6), and SNr (Fig. 7). The proportions of excitatory and inhibitory inputs (Supplementary Table 1) together with the parameters of the model neurons (Supplementary Table 2) generated baseline (DBS-OFF) firing rates which corresponded to *in vivo* recordings. The left side of Fig. 5 shows the simulated membrane potential (accounting also for action potential generation) before and during stimulation across a range of frequencies for Vim, whereas the right side shows an exemplary *in vivo* Vim neuron. A selective oscillation due to stimulus entrainment is reproduced by the model neuron for DBS at 20Hz. The model neuron can moreover partially reproduce the transient excitatory responses at DBS onset with 50Hz and 100Hz stimulation and 30Hz to some degree; however the transient excitatory responses within the model are of shorter latency. For STN (Fig. 6), the simulated (left) neuronal firing compared to baseline decreases for DBS at ≥30Hz, corroborating experimental data (exemplary *in vivo* STN neuron portrayed on the right side). Neuronal firing rates are substantially attenuated with DBS at 50Hz and 100Hz (as is the case experimentally) due to non-selective synaptic depression. For SNr (Fig. 7), the simulated (left) neuronal firing rate decreases dramatically beginning at 20Hz due to the dominant inhibitory presynaptic currents, corroborating experimental data (exemplary *in vivo* SNr neuron portrayed on the right side). The model neuron fails to generate action potentials for DBS ≥50Hz (as is the case experimentally) due to non-selective synaptic depression. Time-domain histograms are also presented in each figure (Fig. 5, 6, 7) which were generated by averaging the neuronal firing rates of 10 modelled neurons for each respective structure across 2s of stimulation at each frequency.

**Fig. 5.**
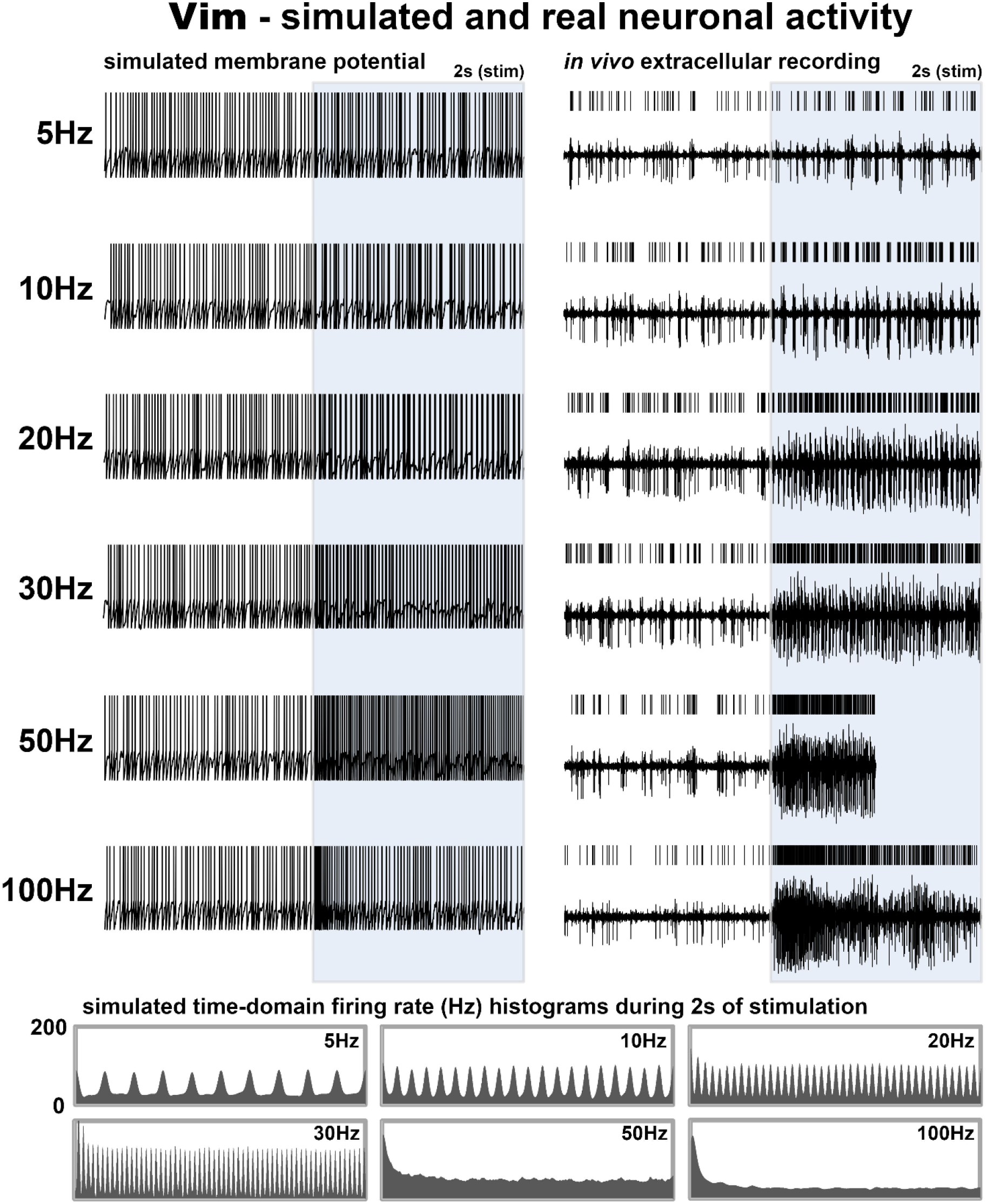
Computational time-domain membrane potential for Vim. The left panels show the membrane potential of a model Vim neuron immediately before (non-shaded) and during (shaded) DBS across a range of frequencies. The right panels are exemplary recordings from an in vivo human Vim neuron (stimulation for 50Hz was limited to 1s). The bottom-most panels are time-domain firing rate histograms generated by averaging across 10 model Vim neurons. A selective oscillation due to stimulus entrainment is reproduced by the model neuron for DBS at 20Hz. The model neuron can partially generate the transient excitatory responses at DBS onset with 50Hz and 100Hz stimulation and 30Hz to some degree; however, the transient excitatory responses within the model are of shorter latency.

**Fig. 6.**
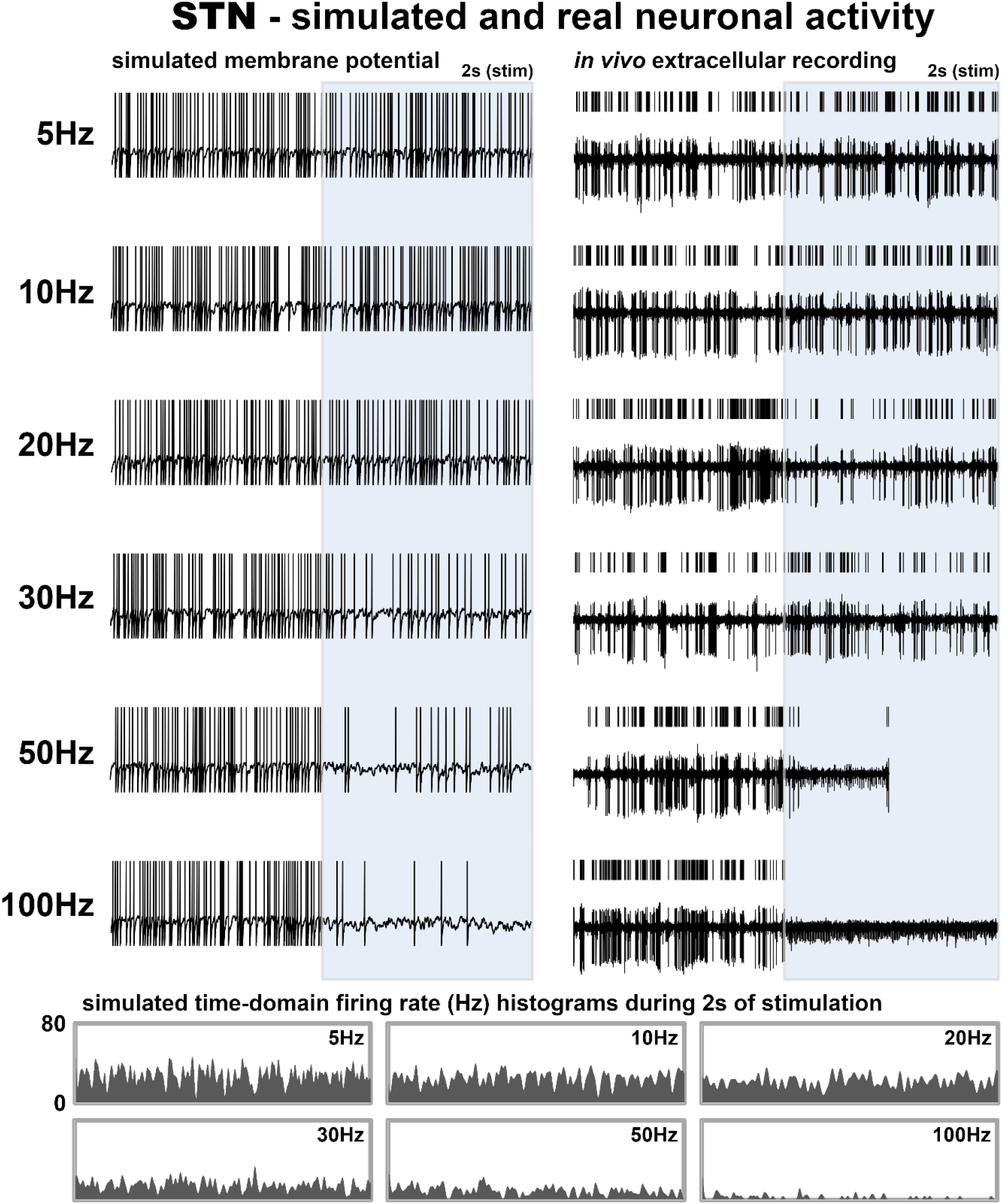
Computational time-domain membrane potential for STN. The left panels show the membrane potential of a model STN neuron immediately before (non-shaded) and during (shaded) DBS across a range of frequencies. The right panels are exemplary recordings from an in vivo human STN neuron (stimulation for 50Hz was limited to 1s). The bottom-most panels are time-domain firing rate histograms generated by averaging across 10 model STN neurons. The neuronal firing rate compared to baseline decreases for DBS at ≥30Hz, corroborating experimental data. The modelled neuronal firing rates are substantially attenuated with 50Hz and 100Hz (as is the case experimentally) due to non-selective synaptic depression.

**Fig. 7.**
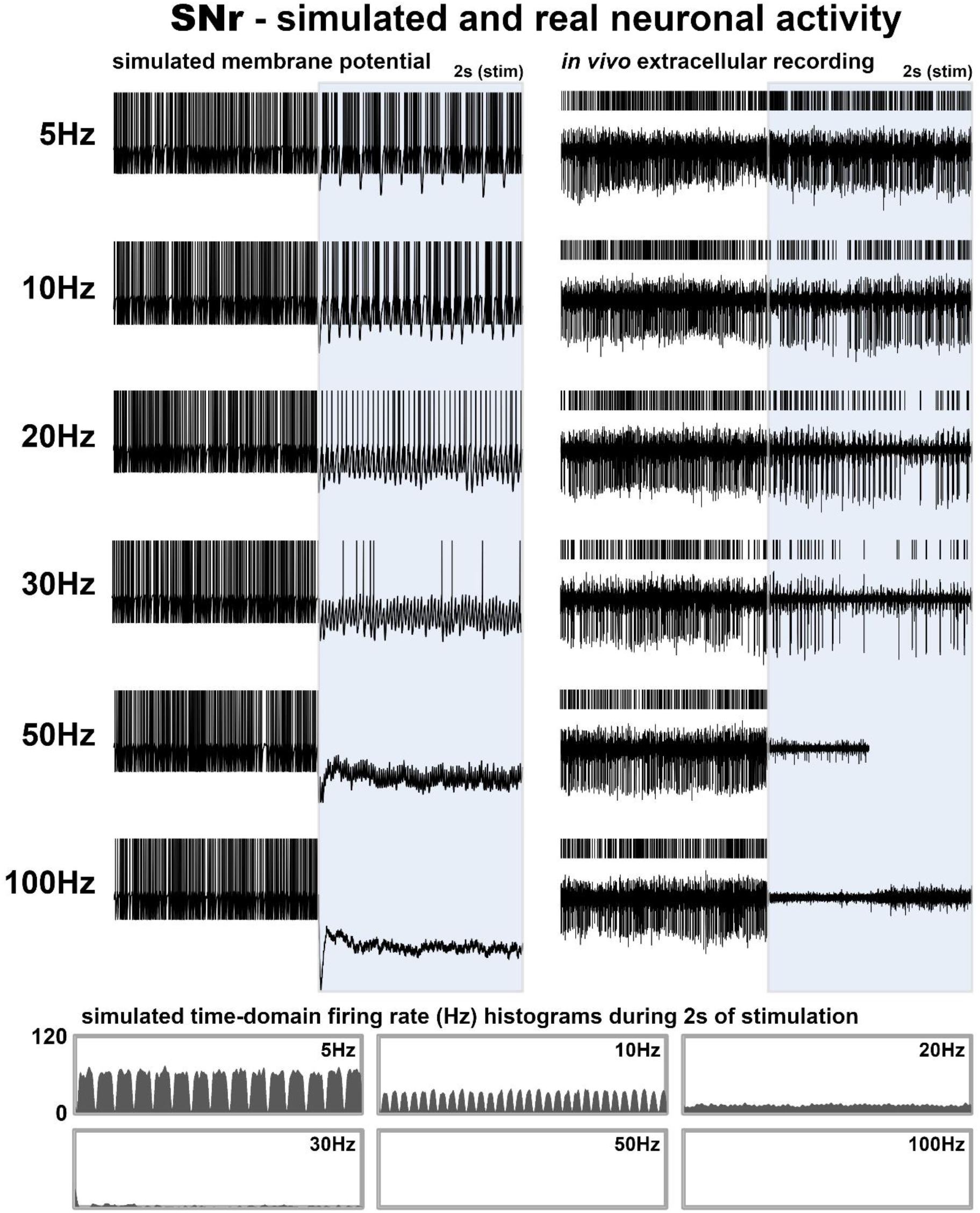
Computational time-domain membrane potential for SNr. The left panels show the membrane potential of a model SNr neuron immediately before (non-shaded) and during (shaded) DBS across a range of frequencies. The right panels are exemplary recordings from an in vivo human SNr neuron (stimulation for 50Hz was limited to 1s). The bottom-most panels are time-domain firing rate histograms generated by averaging across 10 model SNr neurons. Due to the dominant inhibitory presynaptic currents, the neuronal firing rate decreases dramatically beginning at 20Hz, corroborating experimental data. The model neuron fails to generate action potentials for DBS ≥50Hz (as is the case experimentally) due to non-selective synaptic depression

## Discussion

### Site-specific and frequency-dependent stimulation effects

At the somatic level, electrical stimulation is both site-specific and frequency-dependent. In Vim and Rt, neuronal activity could be upregulated, whereas in STN and SNr it was downregulated. These mechanistic disparities across brain regions are most likely explained by anatomical differences in local microcircuitries, in that the effects appeared dependent upon the relative distributions of excitatory and inhibitory inputs converging at target neurons (Chiken and Nambu, 2014). The experimental findings demonstrated that neuronal activity in any brain region could be suppressed either selectively in regions with a high predominance of inhibitory inputs or non-selectively if high enough stimulation frequencies were used. Neuronal excitation, however, could only be achieved when electrical stimulation was delivered to brain regions with a high predominance of glutamatergic inputs. While these bimodal effects (excitatory vs. inhibitory) with low stimulation frequencies were likely attributable to presynaptic activation, the loss of site-specificity and convergence towards neuronal suppression with sustained HFS (≥100Hz) was most likely attributable to synaptic depression (Liu et al., 2012; Milosevic et al., 2018a; Rosenbaum et al., 2014; Steiner et al., 2019). This phenomenon of short-term synaptic plasticity can be defined as a reversible decrease in synaptic efficacy, caused by the depletion of readily releasable neurotransmitter vesicle pools when successive stimuli are delivered at a fast ratel; a reduction of presynaptic calcium conductance; and/or the inactivation of neurotransmitter release sites due to delayed recovery from vesicle fusion events (Dittman and Regehr, 1998; Fioravante and Regehr, 2011; Rizzoli and Betz, 2005; Rosenmund and Stevens, 1996; Zucker and Regehr, 2002).

Our computational model was designed to test our two main hypotheses, (i) that the post-synaptic responses (i.e. neuronal output) to single pulses of electrical stimulation were mediated by the proportions of inhibitory vs. excitatory inputs to the stimulated neuron, and (ii) that weakened synaptic transmission fidelity over time with higher stimulation frequencies was mediated by short-term synaptic plasticity. As such, the biophysical modelling approach takes into consideration both anatomical (local microcircuitry) and physiological (short-term synaptic dynamics) properties. At stimulation frequencies below the threshold for synaptic depression (i.e. <20-30Hz) (Liu et al., 2012; Milosevic et al., 2018a), our model showed that neuronal responses were the result of a temporal summation of stimulus-evoked responses. In structures with predominantly excitatory inputs, this led to increases in neuronal output, whereas the opposite occurred in structures with predominantly inhibitory inputs. Beyond the threshold for synaptic depression, the strengths of successive stimulus-evoked responses were progressively reduced (i.e. a loss of synaptic transmission fidelity). In the Vim, with high frequencies, we observed an initial excitatory response which weakened over time. The SNr too was affected by synaptic depression, although the impact upon neuronal firing was unable to be observed since the stimulus-evoked inhibitory responses are so strong that neuronal silencing can be achieved by temporal summation of inhibitory events alone (thus one cannot observe the neuronal output). However, we have previously shown progressive, frequency-dependent decreases to the amplitudes of extracellular evoked field potentials in SNr with stimulation frequencies ≥20Hz (Milosevic et al., 2018a). It may then be assumed that since synaptic depression would weaken the strength of inhibitory synaptic transmission, neuronal firing should increase via disinhibition. However, our model shows non-selective synaptic depression of both inhibitory and excitatory synaptic currents. This is indeed supported by experimental work in rodent STN slices which demonstrated that pharmacologically-isolated excitatory and inhibitory postsynaptic potentials were both depressed during HFS (Steiner et al., 2019). High-frequency DBS has therefore been considered a “functional deafferentation” (Anderson et al., 2004). This would also explain the suppression of somatic firing in STN with higher stimulation frequencies, whereas the stimulus-evoked responses with lower frequencies produced rather weak net inhibitory responses due to the more homogenous distribution of excitatory and inhibitory inputs to STN.

### Stimulation of the basal ganglia

Individual stimulation pulses elicited net inhibitory responses in both STN and SNr. These responses are corroborated by anatomical studies which suggest a predominance of GABAergic inputs to each of these structures (summarized in the STAR Methods subsection “Computational: Presynaptic inputs”). The greater predominance of inhibitory inputs to SNr thus explains the stronger inhibitory responses compared to STN. While single pulses of stimulation at low frequencies elicited inhibitory responses, with higher stimulation frequencies (i.e. ≥20Hz), neuronal inhibition/suppression was even more easily achieved likely due to a combined effect of inhibitory temporal summation and non-selective synaptic depression.

The canonical model of Parkinson’s disease suggests that the activity of the STN is pathologically increased as a result of a lack of downregulation of D2-dopamine-receptor-mediated “indirect-pathway” projections (Albin et al., 1989; Alexander and Crutcher, 1990; DeLong, 1990; Surmeier et al., 2007). The subsequent increased excitatory drive of subthalamo-pallidal and nigral projections, in combination with a lack of upregulation of D1-dopamine-receptor-mediated “direct-pathway” projections, leads to a pathological overactivity of the GPi and SNr. As such, in Parkinson’s disease, the STN, SNr, and GPi may all benefit from the suppression of neuronal overactivity, and/or stimulation-induced abolishment of pathophysiological oscillations (Kühn et al., 2008) which manifest throughout the disease process (Brown, 2003). It must also, however, be considered that while somatic firing would be suppressed in these structures, each DBS pulse may initiate an action potential along the efferent axons (McIntyre et al., 2004). Thus, high-frequency DBS may work to replace pathological somatic firing with regularized high-frequency output.

### Stimulation of the thalamic ventral intermediate nucleus

In the Vim, individual stimulation pulses elicited time-locked excitatory responses. While stimulus-evoked inhibitory responses in SNr and STN were likely caused by the predominance of inhibitory afferent inputs, the excitatory responses elicited in Vim were most likely explained by the predominant excitatory afferent inputs. In high-frequency stimulation of the Vim, somatic firing was transiently upregulated at the start of stimulation, but this effect weakened quickly and progressively over time. As our model demonstrates, the initial excitatory response can likely be explained by synaptic activation of the predominant glutamatergic afferents, while the subsequent weakening is likely the result of synaptic depression (Ran et al., 2009) due to the fast rate at which subsequent stimuli are delivered. In an intracellular sensorimotor thalamic rat brain slice, it was indeed shown that 125Hz stimulation resulted in an initial depolarization response coupled with increased neuronal firing, followed quickly by (complete or incomplete) repolarization coupled with suppression of neuronal firing (Anderson et al., 2004). These responses were reversed by application of glutamate receptor and calcium channel antagonists, demonstrating their presynaptic nature. The neuronal suppression which occurred after the initial excitation was indeed suggested to be the result of synaptic depression.

The pathology of essential tremor is not well understood, but is increasingly being studied, with several lines of evidence indicating a dysfunction and possible degeneration of the cerebellum (Louis et al., 2007, 2019); which subsequently projects to the Vim. Postmortem literature has revealed pathologies including but not limited to Purkinje cell loss (Axelrad et al., 2008; Shill et al., 2008), Purkinje cell axonal and dendritic swelling (Louis et al., 2011; Yu et al., 2012), and reduced cerebellar GABAergic tone (Paris-Robidas et al., 2012). These pathological changes may indeed drive the characteristic neurophysiological changes that occur in patients with essential tremor (Helmich et al., 2013), including tremor-related oscillations at the local field potential (Kane et al., 2009) and single-unit (Lenz et al., 1988) levels which are coherent with tremor at the periphery. As such, the therapeutic efficacy of high-frequency Vim-DBS might be explained by suppression of neuronal firing and/or abolishment of pathophysiological tremor-related oscillatory activity (Milosevic et al., 2018b). Conversely, low-frequency stimulation-induced tremor (Barnikol et al., 2008; Swan et al., 2016) might explained by periodic neuronal entrainment which mimics tremor-related neuronal activity.

### Stimulation of the reticular thalamus

In the Rt, like in the Vim, individual stimulation pulses elicited time-locked excitatory responses. Also like in the Vim, the upregulation of neuronal firing during stimulation at ≥20Hz weakened over time, and with 100Hz stimulation, the excitatory responses were limited to the first ~0.5s of the stimulation train, after which somatic firing was reduced to or below baseline. A recent *in vivo* nonhuman primate study (Lymer et al., 2019) demonstrated inhibitory responses which followed 0.5s trains of microstimulation at 100Hz (note: neuronal firing was not assessed during stimulation) hypothesized to have been mediated by GABAergic afferent inputs to the Rt. Here, we showed that 0.5s 100Hz stimulation trains in fact initially drove neuronal firing during stimulation, which likely reflects the prominent glutamatergic input, after which neuronal firing was suppressed likely due to synaptic depression. Slice work has indeed demonstrated that electrical stimulation of the Rt can elicit EPSCs (Ulrich and Huguenard, 1995) and neuronal spiking (Landisman et al., 2002; Schlag and Waszak, 1971). Moreover, stimulation of layer VI cortical neurons (Gentet and Ulrich, 2004), and ventrobasal (Gentet and Ulrich, 2003) and ventrolateral (Bazhenov et al., 1999) thalamic neurons elicited excitatory responses in Rt.

A conventional application of Rt-DBS does not exist; however, in animal studies, Rt stimulation has been implicated as a possible treatment for various forms of epilepsy (Chang et al., 2017; Nanobashvili et al., 2003, 2012; Pantoja-Jiménez et al., 2014; Wang and Wang, 2017). Hypersynchronization of Rt output has been implicated in spike-wave seizures (Blumenfeld, 2005), supporting a rationale for Rt-DBS for intractable epilepsy (Krauss and Koubeissi, 2007). Moreover, the GABAergic output of the Rt has an important role in controlling excitability throughout the thalamus as it is known to project to many different thalamic nuclei (Cox et al., 1997; Kim et al., 1997; Kultas-Ilinsky et al., 1995; Steriade et al., 1984). In experimental applications of Rt-DBS, it must be considered that upregulation of Rt neuronal activity would have a widespread inhibitory effect on downstream thalamic structures, while suppression of Rt activity would lead to disinhibition (unless its output is indeed replaced by regularized DBS pulses). While not specifically applied to Rt, DBS of other nodes of the ascending reticular activating system have also been investigated for minimally conscious state (Chudy et al., 2018; Yamamoto and Katayama, 2005) due to the role of this system in behavioural arousal and consciousness. Moreover, stimulation of the pontomesencephalic tegmentum area has been shown to modulate sleep in patients with Parkinson’s disease (Arnulf et al., 2010; Lim et al., 2009). Considering the role of Rt in arousal and attention (Lewis et al., 2015; McAlonan et al., 2008) and the role of synchronous Rt neuronal activity (which can be induced by stimulation as seen in Supplementary Fig. 2) during sleep (Landisman et al., 2002), stimulation of the Rt may have a possible role in sleep disorders.

### Translational implications

The selectively bimodal and frequency-dependent somatic responses described here should be taken into consideration in the development of novel stimulation paradigms and DBS indications. In applications of DBS which utilize a high stimulation frequency, suppression of somatic firing is likely achieved. Stimulation paradigms which utilize low stimulation frequencies and are applied to areas of the brain with predominantly glutamatergic inputs may depend upon periodic facilitation of somatic firing, with one possible example being low-frequency pedunculopontine-DBS (Moreau et al., 2009). Low-frequency stimulation in an area of the brain with predominantly inhibitory inputs may on the other hand cause periodic inhibitions. In either case, low-frequency stimulation can induce oscillatory neuronal behaviour (as seen in Supplementary Fig. 2). Knowledge of the site-specific and frequency-dependent properties of DBS can inform the development of novel stimulation paradigms such as closed-loop stimulation for on demand upregulation or downregulation of neuronal firing, or for induction or disruption of neuronal oscillations. Indeed, stimulation parameters are often decided upon empirically. Based on the findings presented here, knowledge of the local microcircuitry (distribution of afferent inputs) inherent to the stimulated brain region (i.e. therapeutic targets of interest for DBS application) may allow us to infer/predict the stimulation frequency response properties. As such, our comprehensive computational model may represent a valuable tool for physiologically-informed stimulation programming and paradigm development in prospective DBS targets and indications, particularly as our model was developed based on *in vivo* experimental data from the human brain.

### Considerations and limitations

While suppression of somatic activity may indeed be therapeutic (Bergman et al., 1990; Levy et al., 2001; Wichmann et al., 1994), orthodromic and antidromic axonal effects of electrical stimulation must also be considered (McIntyre and Hahn, 2010). If each DBS pulse generates an axonal action potential (McIntyre et al., 2004), then the overall “neuronal output” should be considered as the summation of the somatic firing rate (which would have been influenced by afferent axon activations) and stimulation frequency (i.e. efferent axon activation); this summation is incorporated into Fig. 1B. Thus, HFS applications which completely suppresses somatic firing would replace neuronal output with regular outputs corresponding to the stimulation frequency. Although we did not record from any structures downstream of the stimulation site, it is perhaps possible to infer the downstream effects based on the results presented here. For example, in stimulation of the STN, activation of the glutamatergic subthalamo-pallidal/nigral efferents may cause excitatory responses downstream (Boulet et al., 2006; Galati et al., 2006; Hashimoto et al., 2003), especially if lower stimulation frequencies are used; whereas with higher stimulation frequencies the downstream glutamatergic drive may in fact be weakened (Maltête et al., 2007; Tai et al., 2003; Zheng et al., 2011) due to synaptic depression. Further studies are warranted in order to better understand the possible orthodromic (and antidromic) network phenomena of DBS (Alhourani et al., 2015). Moreover, studies relating to the downstream and upstream DBS effects would allow us to better understand the mechanisms of DBS applied to white matter tracts (such as forniceal-DBS). Another notable limitation of this study is that the applied stimulation trains were limited to short durations; stimulation effects over longer durations are yet to be validated. Moreover, while this study aimed to elucidate differential mechanisms involved in stimulation of various brain structures, behavioural and clinical correlates were not assessed here directly. However, the high-frequency microstimulation applied to Vim was shown to be effective at suppressing tremor (Milosevic et al., 2018b) confirming that the stimulation parameters used were clinically relevant. Moreover, the stimulation parameters used here (in particular, the 100Hz microstimulation trains) are comparable in terms of total electrical energy delivered during clinically-applied DBS macrostimulation (Milosevic et al., 2019) though are of greater current density due to the smaller stimulating surface. Finally, an important limitation of this work is that the explanation of site-specific mechanistic disparities based on the proportions of inhibitory/excitatory afferent inputs does not account for the possible contributions of glia (Bekar et al., 2008; Campos et al., 2020; Salatino et al., 2017; Tawfik et al., 2010) nor neuromodulatory inputs (Lavian et al., 2018; Lavoie et al., 1989) which should be considered in future work.

### Conclusion

The presented results demonstrate the site-specific and frequency-dependent neuronal effects of extracellular stimulation. Neuronal suppression could be achieved either by stimulus-evoked inhibitory events in structures with predominantly GABAergic inputs (STN and SNr) or non-selectively when sustained HFS was delivered. Stimulus-evoked neuronal excitatory responses were exclusive to structures with predominantly glutamatergic inputs (Vim and Rt), particularly with lower stimulation frequencies. Our computational model showed that the bimodal site-specific stimulus-evoked responses could be explained by differences in the distributions of inhibitory and excitatory inputs to the stimulated target structures, whereas convergence towards neuronal suppression with sustained HFS could be explained by synaptic depression.

## Methods

### Experimental: Patients and neurons

115 neurons from patients with Parkinson’s disease (n=45) or essential tremor (n=14) were included in this study for the various analyses which are detailed below. All experiments conformed to the guidelines set by the Tri-Council Policy on Ethical Conduct for Research Involving Humans and were approved by the University Health Network Research Ethics Board. Moreover, each patient provided written informed consent prior to taking part in the studies.

### Experimental: Protocols

Neurophysiological mapping procedures were performed during awake DBS surgeries (after overnight withdrawal from medications) using two closely spaced microelectrodes (600μm apart, 0.1-0.4MΩ impedances) (Levy et al., 2007). Techniques for identification of neurons of the Rt, STN, SNr (Hutchison et al., 1998), and Vim (Basha et al., 2014; Milosevic et al., 2018b) have been previously reported. While recording from a well-isolated single neuron (in one of the aforementioned structures) using one microelectrode, microstimulation was delivered at different frequencies using the adjacent microelectrode. Recordings were obtained using two Guideline System GS3000 amplifiers (Axon Instruments, Union City, CA) and signals were digitized at ≥12.5kHz with a CED 1401 data acquisition system (Cambridge Electronic Design, Cambridge, UK). Microstimulation was delivered using an isolated constant-current stimulator (Neuro-Amp1A, Axon Instruments, Union City, CA) with symmetric 0.3ms biphasic pulses (cathodal followed by anodal).

To generate stimulation frequency response functions, stimulation trains were delivered at 1Hz (10 pulses), 2Hz (20 pulses), 3Hz (60 pulses), 5Hz (50 pulses), 10Hz (50 pulses), 20Hz (60 pulses), 30Hz (60 pulses), 50Hz (50 pulses), and 100Hz (50 pulses) using 100μA and a 0.3ms biphasic pulse width. This frequency response protocol was executed at 9 Vim (n_patients_=5), 11 Rt (n_patients_=11), 27 STN (n_patients_=16), and 14 SNr (n_patients_=9) recording sites. The average firing rates of all structures during stimulation at each frequency were plotted. The data for STN and SNr were previously collected (Milosevic et al., 2018a), whereas Vim and Rt data for this study were unique. Longer trains (≥3s) of 100Hz stimulation were also delivered to the aforementioned Vim, Rt, and SNr neurons. 100Hz long train data for STN (44 neurons, n_patients_=20) were previously collected (Milosevic et al., 2019), as were a subset of 100Hz and 200Hz Vim (14 recording sites, n_patients_=9) data (Milosevic et al., 2018b).

### Experimental: Offline analyses and statistics

The narrow stimulus artifacts were removed offline (0.5ms from stimulus onset), recordings were high pass filtered (≥300Hz) to better isolate the single units, and template matching was done using a principal component analysis method in Spike2 (Cambridge Electronic Design, UK). For the frequency response protocol (in which ≤60 stimulation pulses were delivered at each frequency), the average pre-stimulation baseline firing rates were measured, as were the firing rates during the various stimulation trains. Kolmogorov-Smirnov tests confirmed normality of data. One-way repeated measures ANOVA tests (stimulation frequency as a within-subject factor) were carried out, and in the event of a significant main effects, Bonferroni-corrected (nine comparisons) post hoc t-tests were used to compare firing rates during the various stimulation trains to the pre-stimulation baseline firing. ANOVA effect sizes (η^2^; proportion of variance associated with the main effects) and t-test effect sizes (Cohen’s d_z_; standardised difference between two means) were also determined. One neuron from the Vim group and one neuron from the Rt group were excluded from statistical analyses due to missing data points (i.e. incomplete stimulation protocol). To investigate the effects of individual stimulation pulses, an average peristimulus histogram (120ms total width, 20ms offset, 2ms bins) was created for all neurons using the 5Hz stimulation trains (50 pulses). The average firing rates during the 20ms pre-stimulus periods were compared to the average firing rates during 20ms and 40ms post-stimulus periods using Bonferroni-corrected (two comparisons) two-tailed paired t-tests, and effect sizes (Cohen’s d_z_) were calculated. Finally, to investigate possible time-varying responses throughout the duration of stimulation trains (rather than average responses to individual stimuli), time-series histograms (2-3s total width, no offset, 50ms bins) were created for 5Hz, 10Hz, 20Hz, 30Hz, and long trains of 100Hz (and 200Hz long trains for Vim) stimulation trains (i.e. trains where ≥2s of stimulation was delivered). The attenuations of excitation over time in Vim and Rt during stimulation trains of ≥20Hz were modelled by double exponential functions. Similar histograms were also created for the shorter trains (≤1s) of stimulation at 50Hz and 100Hz (0.5-1s total width, no offset, 20ms bins); these are available in Supplementary Fig. 1.

### Computational: Model framework

To model the effect of DBS pulses on the afferents of the stimulated nuclei, we used a leaky integrate and fire (LIF) single neuron model, together with a TM model of short-term synaptic plasticity (Tsodyks et al., 1998). Each model neuron received a certain number of excitatory and inhibitory presynaptic inputs whose proportion was obtained using morphological data (ex. (Bottcher, 1975)). In addition to these inputs, the background synaptic activity (Destexhe et al., 2001) was modeled by an Ornstein-Uhlenbeck (OU) process and added to the model neuron to reproduce the impact of synaptic noise that exists *in vivo* (Destexhe et al., 2001, 2003). In accordance with our first hypothesis, each DBS single pulse simultaneously activated all presynaptic inputs (Fig. 3A). This simultaneous activation was modeled by artificially generating precise spike times which correspond to the arrival of each DBS pulse in the presynaptic inputs. We utilized our modeling framework to recreate the neuronal firing in Vim, STN, and SNr in response to stimulation trains with frequencies of 1, 2, 5, 10, 20, 30, 50, 100Hz. Model generation for Rt neurons was omitted due to redundancy as the model parameters are identical to Vim except for the parameter which underlies the baseline firing rate (elaborated upon in the “Computational: Parameter settings” subsection below).

### Computational: Presynaptic inputs

The vast majority of inputs to the Vim are glutamatergic projections from the dentate nucleus of the cerebellum (Asanuma et al., 1983; Ilinsky and Kultas-Ilinsky, 2002; Kultas-Ilinsky and Ilinsky, 1991; Kuramoto et al., 2011) and reciprocal connections from cerebral cortex (Kakei et al., 2001; Stepniewska et al., 1994), with less prominent inputs coming via inhibitory Rt projections (Ambardekar et al., 1999; Ilinsky et al., 1999; Kuramoto et al., 2011). The Rt is a thin sheet of neurons that forms a shell around the lateral and anterior borders of the dorsal, and to some extent ventral thalamus (Jones, 1975). It is primarily innervated by collateral branches of glutamatergic thalamocortical and corticothalamic projections (Crabtree, 1992a, 1992b; Gonzalo-Ruiz and Lieberman, 1995; Jones, 1975; Pinault, 2004), but also receives less prominent GABAergic innervation from the GPe and SNr (Asanuma, 1994; Hazrati and Parent, 1991; Paré et al., 1990); like Vim, the majority of afferent inputs to Rt are glutamatergic. The vast majority (~90%) of inputs to the SNr are GABAergic, projecting from the striatum and globus pallidus externus (GPe) (Bolam et al., 2000; Parent and Hazrati, 1995), whereas the STN receives a more homogenous convergence of GABAergic and glutamatergic inputs from the GPe (Baufreton et al., 2009) and motor cortical areas (Nambu et al., 2002) respectively (Parent and Hazrati, 1995; Rinvik and Ottersen, 1993). While the mixed inputs are more homogenous in STN, electron-microscopy work suggests that GABAergic terminals nevertheless outnumber glutamatergic terminals (Kita and Kita, 2012). Estimates of the proportions of inhibitory and excitatory inputs are provided in Supplementary Table 1.

In the model, an ensemble of 500 neurons, each of which was modeled by an LIF model, produced inputs to the stimulated nuclei. Each neuron receiving a random input (modeled by OU process of time constant 5ms) fired at the rate of about 5Hz (the total average firing rate across neurons is equal to 5±0.7Hz). Given an estimated proportion of excitation and inhibition, these neurons were labeled as excitatory and inhibitory neurons, and their spikes were fed to the stimulated nuclei through the TM model. The total synaptic current was obtained as a linear combination of presynaptic excitatory (I_exc_) and inhibitory currents (I_inh_):

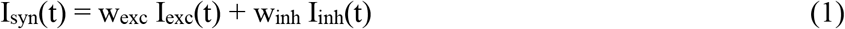

where w_exc_ and w_inh_ denote the weights of excitatory and inhibitory currents, respectively. These weights, together with the mean and standard deviation of the background synaptic current, were tuned to reproduce the neuronal firing rates at the baseline (DBS-OFF) as well as in response to DBS with different frequencies (Supplementary Table 2).

### Computational: Synapse model

We utilized the TM equations to model the function of short-term synaptic plasticity:

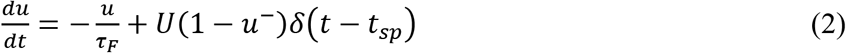

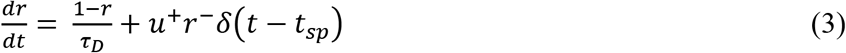

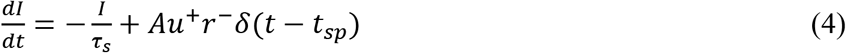

where *u* indicates utilization probability, i.e., the probability of releasing neurotransmitters in synaptic cleft due to calcium ion flux in the presynaptic terminal. Upon the arrival of each presynaptic spike, *t_sp_*, *u* increases by *U*(1 − *u*^−^) and then decays to zero by the facilitation time constant, *τ_f_*. As well, the vesicle depletion process – due to the release of neurotransmitters – was modeled by (2) where *r* denotes the fraction of available resources after neurotransmitter depletion. In contrast to the increase of *u* upon the arrival of each presynaptic spike, *r* drops and then recovers by depression time constant *τ_D_* to its steady state value of 1. The competition between the depression (*τ_D_*) and facilitation (*τ_f_*) time constants determines the dynamics of the synapse. In the TM model, *U*, *τ_f_*, and *τ_D_* are three parameters that determine the type and dynamics of the synapse. In (4), *I* and *τ_s_* indicate the presynaptic current and its time constant, respectively. The time constants of the excitatory and inhibitory inputs are 3ms and 10ms, respectively.

### Computational: Background synaptic activity

Similar to (Destexhe et al., 2001), we used OU process of the time constant of 5ms to represent the effect of synaptic noise. The OU process can be written as:

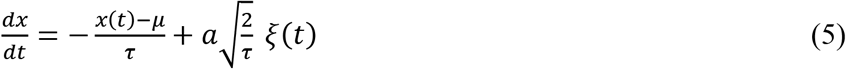

where *ξ* is a random number drawn from a Gaussian distribution with 0 average and unit variance. *τ* is the time constant, *μ* and *α* indicate the mean and standard deviation of variable *x,* respectively.

### Computational: Neuron model

The dynamics of membrane potential in an LIF model can be written as:

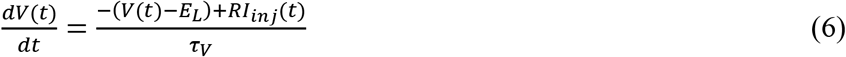

where *E*_L_=−70mV, *R* = 1MΩ, and *τ_V_* = 10ms. *I_inj_* indicates the total injected current to the model neuron (i.e., I_syn_ plus background synaptic noise (5)). Spike occurs when V≥Vth, where *V_th_* = − 40mV and the reset voltage is −90mV with an absolute refractory period of 1 msec.

### Computational: Parameter settings

The proportions of excitatory and inhibitory neurons (Supplementary Table 1), total synaptic current (Supplementary Table 2), parameters of excitatory (Supplementary Table 3) and inhibitory (Supplementary Table 4) synapses, and time constants of membrane dynamics and synaptic currents (Supplementary Table 5) are available in the Supplementary Material. As mentioned, modelling of Rt was omitted due to redundancy as all parameters are identical to Vim except for w_exc_, w_inh_ (but with almost the same ratio) and parameters of background synaptic noise (Supplementary Table 2) which would mediate the difference in baseline firing between Rt and Vim. Parameters for Rt modelling are nevertheless included within the Supplementary Tables.

### Resource availability

Anonymized experimental data is enclosed within Supplementary Data. The Codes for the computational model are available at https://github.com/nsbspl/DBS_Mechanism_Cellular.

## Acknowledgements

The authors thank Tameem Al-Ozzi for assistance in data collection and the patients for their participation.

## Author contributions

1) Research project: A. Conception, B. Organization, C. Execution.

2) Data and statistical analysis: A. Design, B. Execution, C. Review and Critique.

3) Manuscript: A. Writing of the first draft, B. Review and Critique.

LM: 1A, 1B, 1C, 2A, 2B, 3A

SKK: 1B, 1C, 2C, 3B

MH: 1B, 1C, 2C, 3B

AML: 1B, 1C, 2C, 3B

MRP: 1B, 2C, 3B

WDH: 1A, 1B, 1C, 2C, 3B

ML: 1A, 1B, 1C, 2A, 2B, 3A

## Supplementary material information

### Supplementary material

Supplementary Table 1 – Proportions of excitatory and inhibitory presynaptic inputs

Supplementary Table 2 – Parameters for total current fed to the neuron

Supplementary Table 3 – Parameters of excitatory synapses

Supplementary Table 4 – Parameters of inhibitory synapses

Supplementary Table 5 – Time constants of membrane dynamics and synaptic currents

Supplementary Fig. 1 – Time-domain firing histograms for short trains of 50Hz and 100Hz

Supplementary Fig. 2 – Induction of neuronal oscillation with low-frequency stimulation

### Supplementary data

Experimental data files

