## Supplementary material for "A theoretical framework for the site-specific and frequency-dependent neuronal effects of deep brain stimulation"

### Table of contents

**Supplementary Table 1 –** Proportions of excitatory and inhibitory presynaptic inputs 2

**Supplementary Table 2 –** Parameters for total current fed to the neuron 2

**Supplementary Table 3 –** Parameters of excitatory synapses 2

**Supplementary Table 4 –** Parameters of inhibitory synapses 2

**Supplementary Table 5 –** Time constants of membrane dynamics and synaptic currents 2

**Supplementary Fig. 2 –** Induction of neuronal oscillation with low-frequency stimulation 3

**Supplementary Fig. 1** – Time-domain firing histograms for short trains of 50Hz and 100Hz 4

**Supplementary Table 1 –** Proportions of excitatory and inhibitory presynaptic inputs

| | **number of excitatory neurons** | **number of inhibitory neurons** |
|---|---|---|
| **STN** | 225 (45%) | 275 (55%) |
| **SNr** | 50 (10%) | 450 (90%) |
| **Vim / Rt** | 450 (90%) | 50 (10%) |

**Supplementary Table 2 –** Parameters for total current fed to the neuron

| | **mean (pA)** | **st. dev. (pA)** | $w_e$ | $w_i$ |
|---|---|---|---|---|
| **STN** | 32 | 11 | 1.5 | 1 |
| **SNr** | 55 | 10 | 6 | 4 |
| **Vim** | 30 | 45 | 37.5 | 90 |
| **Rt** | 12 | 10 | 4.37 | 11.4 |

**Supplementary Table 3 –** Parameters of excitatory synapses

| | **facilitation** | | | **depression** | | | **pseudo linear** | | |
|---|---|---|---|---|---|---|---|---|---|
| | $\tau_D$ **(ms)** | $\tau_U$ **(ms)** | **U** | $\tau_D$ **(ms)** | $\tau_U$ **(ms)** | **U** | $\tau_D$ **(ms)** | $\tau_U$ **(ms)** | **U** |
| **STN, SNr, Vim / Rt** | 138 | 670 | 0.09 | 671 | 17 | 0.5 | 329 | 326 | 0.29 |

**Supplementary Table 4 –** Parameters of inhibitory synapses

| | **facilitation** | | | **depression** | | | **pseudo linear** | | |
|---|---|---|---|---|---|---|---|---|---|
| | $\tau_D$ **(ms)** | $\tau_U$ **(ms)** | **U** | $\tau_D$ **(ms)** | $\tau_U$ **(ms)** | **U** | $\tau_D$ **(ms)** | $\tau_U$ **(ms)** | **U** |
| **STN, SNr, Vim / Rt** | 45 | 376 | 0.016 | 706 | 21 | 0.25 | 144 | 62 | 0.29 |

**Note**: For inhibitory presynaptic neurons, the ratios of facilitation, depression, and pseudo linear synapses for all three substructures were 0.3, 0.4, and 0.3, respectively. For excitatory presynaptic neurons, the above ratios were used for STN and SNr; however, 0.5, 0.3, and 0.2, respectively, were used for Vim / Rt.

**Supplementary Table 5 –** Time constants of membrane dynamics and synaptic currents

| | $\tau_V$ **(msec)** | $\tau_{Exc}$ **(msec)** | $T_{Inh}$ **(msec)** |
|---|---|---|---|
| **STN** | 12 | 3 | 10 |
| **SNr** | 10 | 3 | 10 |
| **Vim / Rt** | 50 | 5 | 8.5 |

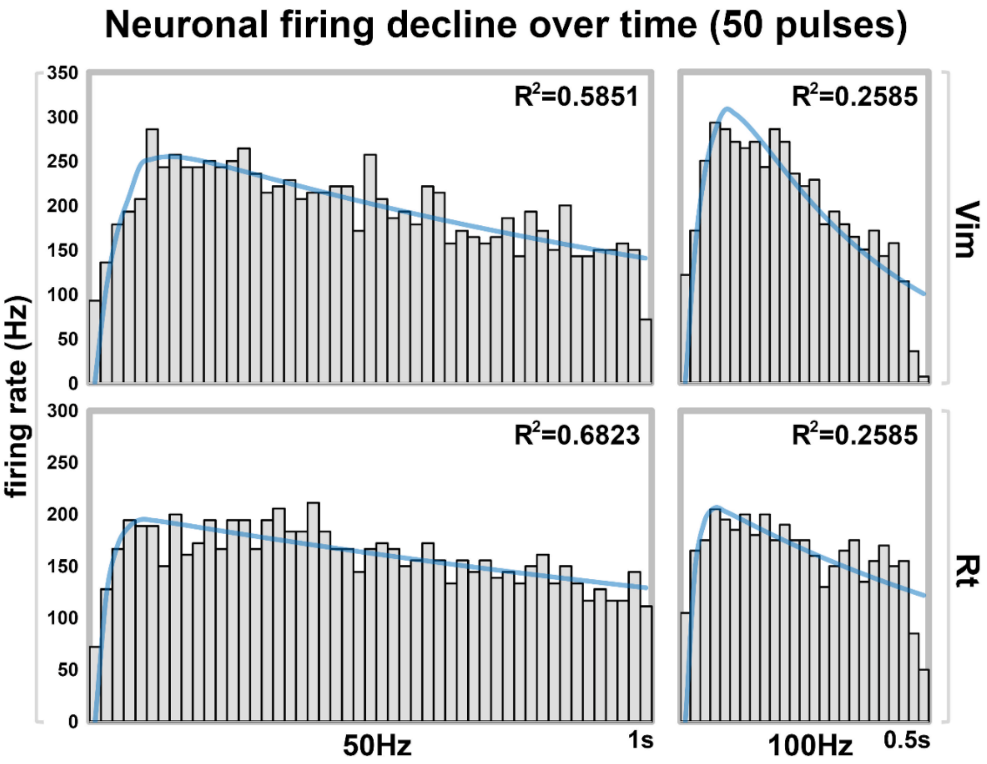


**Supplementary Fig. 1 – Time-domain firing histograms for short trains of 50Hz and 100Hz.** In Vim and Rt, decay of neuronal excitation also occurred with short trains of 50Hz and 100Hz stimulation (50 pulses each).

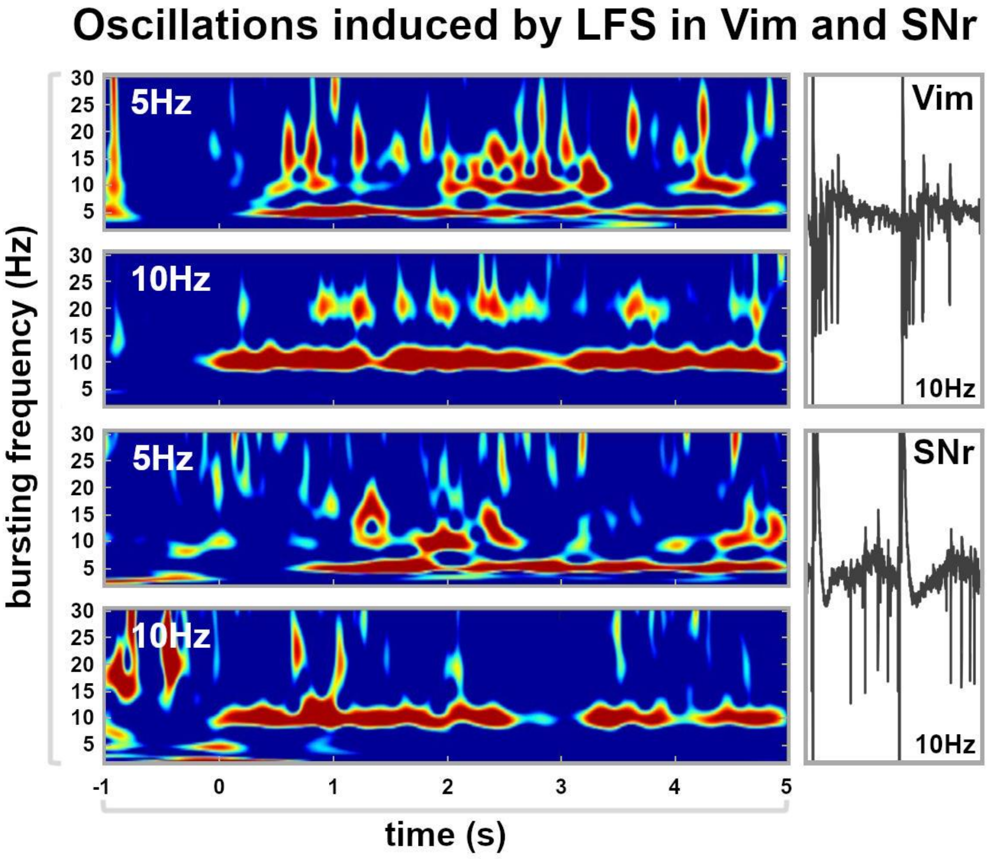


**Supplementary Fig. 2 – Induction of neuronal oscillation with low-frequency stimulation.** In Vim, oscillatory bursting activity was induced (i.e. entrainment) with low-frequency stimulation by way of periodic stimulus-evoked neuronal excitations. In SNr, oscillatory bursting activity was also induced, however, it was caused by periodic stimulus-evoked neuronal inhibitory responses of the otherwise high spontaneous firing rates.
